# Network-based estimation of therapeutic efficacy and adverse reaction potential for prioritisation of anti-cancer drug combinations

**DOI:** 10.1101/2024.09.17.613439

**Authors:** Arindam Ghosh, Vittorio Fortino

## Abstract

Drug combinations, although a key therapeutic agent against cancer, are yet to reach their full applicability potential due to the challenges involved in the identification of effective and safe drug pairs. *In vitro* or *in vivo* screening would have been the optimal approach if combinatorial explosion was not an issue. *In silico* methods, on the other hand, can enable rapid screening of drug pairs to prioritise for experimental validation. Here we present a novel network medicine approach that systematically models the proximity of drug targets to disease-associated genes and adverse effect-associated genes, through the combination of network propagation algorithm and gene set enrichment analysis. The proposed approach is applied in the context of identifying effective drug combinations for cancer treatment starting from a training set of drug combinations curated from DrugComb and DrugBank databases. We observed that effective drug combinations usually enrich disease-related gene sets while adverse drug combinations enrich adverse-effect gene sets. We use this observation to systematically train classifiers distinguishing drug combinations with higher therapeutic effects and no known adverse reaction from combinations with lower therapeutic effects and potential adverse reactions in six cancer types. The approach is tested and validated using drug combinations curated from *in vitro* screening data and clinical reports. Trained classification models are also used to identify novel potential anti-cancer drug combinations for experimental validation. We believe our framework would be a key addition to the anti-cancer drug combination identification pipeline by enabling rapid yet robust estimation of therapeutic efficacy or adverse reaction potential.

## 1. INTRODUCTION

Over the last decades, combination therapy has been crucial in cancer treatments (Hanahan, 2014; Jin et al., 2023). Their advantage is that they can simultaneously target multiple disease-causing pathways while also reducing individual drug doses (Al-Lazikani et al., 2012; Madani Tonekaboni et al., 2018; Zimmermann et al., 2007). Though the use of lower doses can limit dose-related toxicities, the use of two or more drugs simultaneously can lead to adverse drug-drug interactions (DDI) (Blower et al., 2005). Thus, it requires that novel combinations, even involving already approved cancer drugs, are re-screened *in vitro* and/or *in vivo* before actual use in the clinic. However, screening all possible combinations is a combinatorial challenge, where the number of experiments required increases exponentially with the number of drugs. Such exhaustive testing is often infeasible (Poleksic and Xie, 2019; Sun et al., 2013), leading researchers to prioritize the most promising combinations based on *in silico* tools. Computational tools can screen a large number of drug combinations in a relatively shorter time and prioritize a small number of candidate drug pairs for further pre-clinical and clinical trials (Madani Tonekaboni et al., 2018; Rintala et al., 2022). However, a major challenge is developing a robust system that can accurately identify effective and safe drug combinations.

Numerous *in silico* tools, currently available, primarily focus on identifying synergistic or antagonistic drug combinations. They may include monotherapy data, drug-induced gene expression profiles, drug-drug similarity etc. individually or in combination for prediction (Goswami et al., 2015; Huang et al., 2014; Jin et al., 2024; Li et al., 2018; Yang et al., 2015). While they have been successful in identifying drug pairs that will perform better than the individual drugs alone, existsing *in silico* tools do not completely negate the chances of adverse drug-drug interactions (DDI). To address drug combinations with adverse DDI, a separate class of tools is available to predict these interactions (Cheng and Zhao, 2014; Huang et al., 2013; Kastrin et al., 2018). Effectively, this makes the identification of effective and safe anti-cancer drug combinations a two-stage problem. Of late there have been attempts to address the two issues as a single task. Especially, advanced machine learning and deep learning-based methods have demonstrated significant success (Li et al., 2015; Wang et al., 2023; Yue et al., 2023). However, a key limitation has been the amount of data required to train robust models for cancer-type-specific drug combination identification. Another potential disadvantage of such advanced models is their explainability i.e., they often fail to model the relationships of the drug combination targets to the known disease and adverse events associated genes and then use the information to establish exact rules that clearly distinguish effective and adverse drug combinations.

Network-based data mining algorithms using protein-protein interaction (PPI) networks that cover known drug targets, genes/proteins dysregulated by the disease, and those associated with adverse drug reactions could provide an overall solution to the problem (Rintala et al., 2022). Network-based approaches have been successfully used to develop *in silico* strategies for the identification of effective and safe drug combinations. For example, Wu et al., 2010 proposed a novel score to quantify the efficacy and safety of drugs or their combinations based on the expression of disease-related genes and essential genes within the sub-networks affected by drugs or their combinations. On a limited number of Type 2 Diabetes mellitus drug combinations, they showed that an effective drug combination has a higher score than its constituent drugs. While this approach effectively integrates transcriptomic data with known biomolecular interactions, it restricts the identification of drug combinations to those for which the gene expression of the individual drugs is available. To alleviate this bottleneck, (Chen et al., 2016) utilised the drug combination targets to identify the afflicted pathways and devised a synergy score that considers the topological associations between them in a pathway-pathway interaction network. They concluded that synergistic drug combinations are formed by drugs that either act on the same pathway through different targets or regulate a small number of highly connected pathways. A key limitation here is the reliance on known pathways and their interconnectedness for the evaluation of synergy scores. In a similar drug target-based approach, instead of the connections between the pathways, (Cheng et al., 2019) used shortest paths between the drug combination targets and the disease-related genes to calculate a separation distance that can effectively identify disease-specific clinically efficacious drug combinations. According to their findings, for drug combinations to be effective, not only should the drugs have distinct sets of genes/proteins as targets, but they should also target the same disease module. This approach potentially limits itself by using the shortest path to gauge proximity between two gene sets, thereby neglecting significant indirect interactions and broader network context (Tong et al., 2006). Moreover, a single metric used to measure the distance between known drug targets and disease genes is accounted for distinguishing both safe and unsafe drug combinations. While this disease-centric approach might be sufficient to account for therapeutic efficacy it undermines the role of genes/proteins that are linked to adverse effects. This limitation suggests the need to use both disease-related genes/pathways and adverse-effect-related genes/pathways to comprehensively evaluate the efficacy and safety of drug combinations, ensuring that proximity to therapeutic targets does not overshadow potential risks associated with adverse reactions.

This study presents a novel computational approach that integrates network propagation algorithms, like Random Walk with Restart (RWR) (Tong et al., 2006), with gene set enrichment analysis to quantify the potential therapeutic efficacy and adverse-effect-related safety of drug combinations. This integration goes beyond simply considering the shortest paths between drug targets and disease/adverse-effect genes/pathways. RWR leverages the network structure to comprehensively explore and rank intermediate genes (potentially involved in the drug’s mechanism) connecting drug targets to disease or adverse effect genes. It thus provides a more robust ranking of network genes compared to traditional shortest-path algorithms. Furthermore, the incorporation of gene set enrichment analysis reveals if the ranked genes are enriched within disease gene sets, adverse effect gene sets, or potentially both. Subsequently, the enrichment scores could be used as predictors to identify drug combinations that are effective against the disease while posing a lower risk of adverse effects. We systematically evaluate our *in silico* approach by leveraging existing data on cancer cell lines (Zagidullin et al., 2019) and reported adverse DDI from DrugBank (Wishart et al., 2018) to classify drug combinations as "effective" (higher therapeutic efficacy with no adverse drug reaction) or "adverse" (lower therapeutic efficacy with potential adverse drug reaction). Initial assessment through statistical method revealed that effective drug combinations had a higher enrichment of disease-related gene sets, whereas adverse drug combinations had a higher enrichment of adverse-effect-related gene sets. This observation was leveraged to develop predictive systems for identifying and prioritizing effective and safe drug combinations across six cancer types followed by validation using known labelled drug combinations. Additionally, these predictive systems were used to identify novel anti-cancer drug combinations suitable for further experimental validation.

## 2. MATERIALS AND METHODS

### 2.1. Summary of the proposed computational workflow

The framework uses drug target information to derive a surrogate estimate of the potential of the drug combinations to induce therapeutic efficacy or cause adverse effects. RWR, a network propagation algorithm, is first used to identify the key genes/proteins within a PPI network affected by the drug combinations. Subsequently, fast gene set enrichment analysis (FGSEA) is applied to verify whether these genes/proteins are enriched in disease-related gene sets, adverse-effect-related gene sets, or both. The normalised enrichment scores (NES) from FGSEA, alternately referred to as efficacy (for enrichment against disease-related gene sets) or safety (for enrichment against adverse-effect-related gene sets) estimates, are used to compare effective and adverse drug combinations. We define effective drug combinations as those that have potentially higher efficacy with no reported adverse DDI while adverse drug combinations as those that have lower therapeutic efficacy with known adverse DDI. The ability of these estimates to identify potential new drug combinations is assessed using several datasets of labelled drug pairs for treating different cancer types. These datasets are derived from drug combination screening data in cancer cell lines assessing synergism or antagonism (Zagidullin et al., 2019), clinical reports of adverse reaction-inducing DDI from the DrugBank (Wishart et al., 2018) and clinical trial results from CDCDB (Shtar et al., 2022). We compare our approach to an established framework of using direct proximity-based separation distance for prioritising effective drug combinations. Finally, we use our approach to develop predictive systems and identify novel anti-cancer drug combinations using licensed anti-cancer drugs. Figure 1 graphically illustrates this proposed network medicine-based computational framework to rank and identify novel drug combinations for cancer treatments.

**Figure 1:**
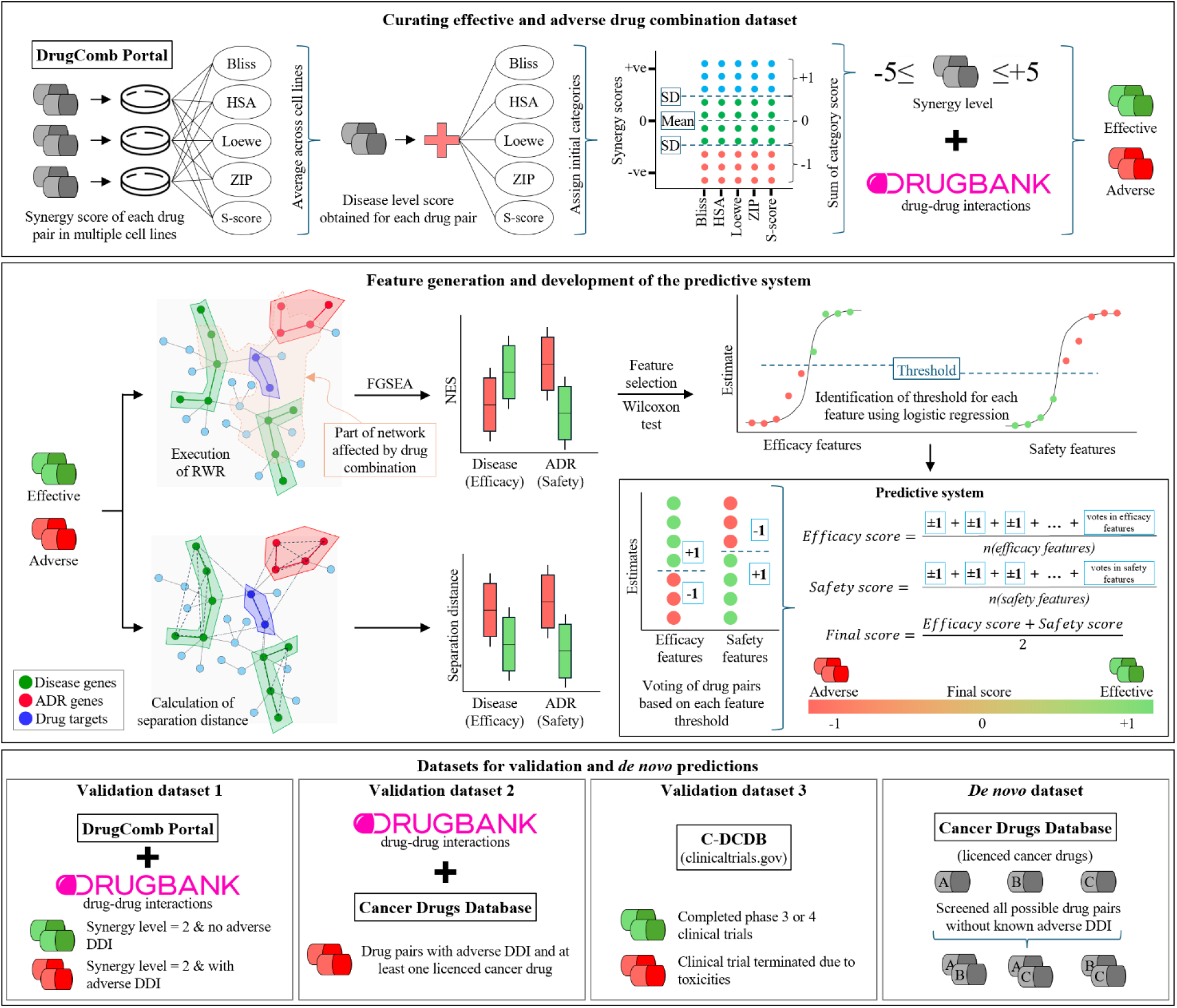
Overview of the network-based pipeline for identification of effective and adverse drug combinations. The pipeline can be divided into three parts. The first part curates a robust dataset of effective and adverse drug combinations. This is done by systematically combining five different synergistic scores of drug combinations in cancer cell lines to obtain a single metric called synergy level that measures the confidence of the combinations to be synergistic or antagonistic. The synergy level is then integrated with reports of cancer-relevant adverse DDI to identify a smaller set of drug combinations labelled as effective or adverse. In the second part, the drug combination targets are used as seed nodes to execute an RWR and identify the zone of the PPI network influenced by them. The list of influenced genes, ranked by their proximity to the target, is then used as input in FGSEA to measure their significance with respect to predefined efficacy and safety gene sets. The resultant NES, alternately referred to as estimates, is used to develop a hierarchical threshold-based predictive system for classifying effective and adverse drug combinations. In this part, the drug combination targets are also used to measure their proximity to efficacy and safety gene sets to verify if the RWR- FGSEA-based efficacy and safety estimates would outperform the proximity-based baseline method in distinguishing effective from adverse drug combinations. The final part of the workflow explores the generalizability of the developed predictive system using external datasets and also identifies novel drug combinations for the treatment of six different cancers.

**Figure 2:**
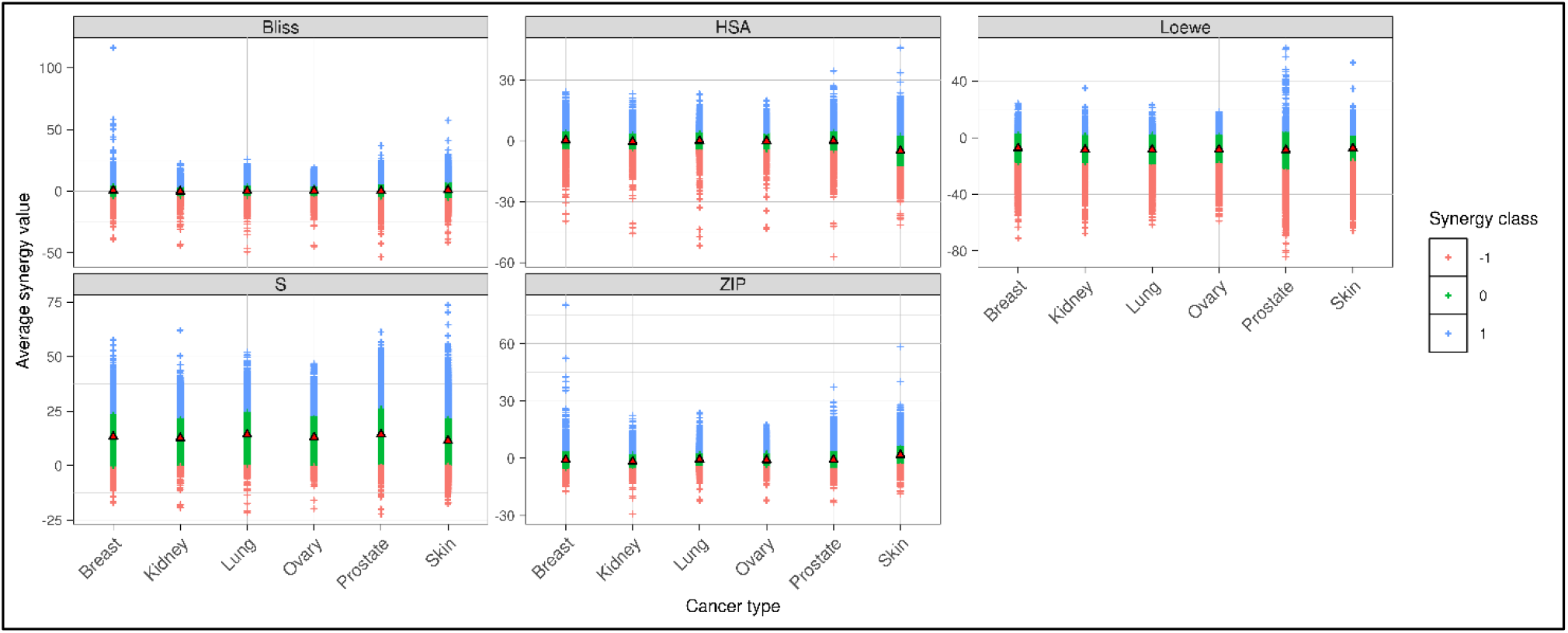
Classification of drug combinations from the DrugComb portal into synergistic (+1), additive (0) and antagonistic (-1) types. For the drug combinations in the portal, the average synergy score across multiple cell lines of the same cancer type was computed followed by calculating the expected mean synergy score for each cancer type. This expected mean synergy score and the corresponding standard deviation within each score type was used as the basis for classification. The drug combinations with synergy scores above 0 and greater than the mean increased by one standard deviation were assigned as synergistic. Alternately, the drug combinations with synergy scores below 0 and less than the mean decreased by one standard deviation were assigned as antagonistic. The drug combinations that did not fall within this criterion were assigned as additive.

### 2.2. Labelling anti-cancer drug combinations using drug combination screening data and clinical reports of drug-drug interactions

In the absence of a gold-standard dataset, effective and adverse drug combinations were defined using *in vitro* drug combination screening data on cancer cell lines and reported adverse reaction-inducing DDI. An overall synergistic score was systematically computed from five different synergy scores – namely Bliss, highest single agent (HSA), Loewe, zero interaction potency (ZIP) and S-Score – of drug combinations in cancer cell lines. These scores were obtained from the FIMM DrugComb portal (Zagidullin et al., 2019) which collects data from several independent drug combination screening studies and provides standardised and harmonised synergy scores for each drug pair in each cell line. The cell lines were mapped to their respective diseases using the NCI thesaurus and those for cancer were retained. Then, for each scoring model, the average synergy score for each drug pair across multiple cell lines of the same cancer type was computed followed by calculating the expected mean synergy score for each cancer type. This expected mean synergy score was used as a baseline of synergy to classify drug combinations as synergistic, additive or antagonistic represented by +1, 0 and −1 respectively (**Figure 1**: Overview of the network-based pipeline for identification of effective and adverse drug combinations. The pipeline can be divided into three parts. The first part curates a robust dataset of effective and adverse drug combinations. This is done by systematically combining five different synergistic scores of drug combinations in cancer cell lines to obtain a single metric called synergy level that measures the confidence of the combinations to be synergistic or antagonistic. The synergy level is then integrated with reports of cancer-relevant adverse DDI to identify a smaller set of drug combinations labelled as effective or adverse. In the second part, the drug combination targets are used as seed nodes to execute an RWR and identify the zone of the PPI network influenced by them. The list of influenced genes, ranked by their proximity to the target, is then used as input in FGSEA to measure their significance with respect to predefined efficacy and safety gene sets. The resultant NES, alternately referred to as estimates, is used to develop a hierarchical threshold-based predictive system for classifying effective and adverse drug combinations. In this part, the drug combination targets are also used to measure their proximity to efficacy and safety gene sets to verify if the RWR- FGSEA-based efficacy and safety estimates would outperform the proximity-based baseline method in distinguishing effective from adverse drug combinations. The final part of the workflow explores the generalizability of the developed predictive system using external datasets and also identifies novel drug combinations for the treatment of six different cancers.). Precisely, drug pairs with synergy scores above 0 and greater than the mean plus one standard deviation were assigned a score of +1. Conversely, those with synergy scores below 0 and lower than the mean minus one standard deviation were assigned a score of −1. The drug pairs that do not fall within this criterion were assigned a score of 0. This new discretised score from five different scoring models was summed to obtain a synergy level for the drug pairs. The synergy level, which can range between -5 to +5, reflects the confidence of the drug combination to be antagonistic or synergistic respectively. Parallely, DrugBank (version 5.1.10) (Wishart et al., 2018) was systematically mined to identify adverse DDI events observed in cancer treatments. Supplementary File 1 lists the DDI events generally associated with all cancer treatments and those specifically associated with treatments of specific cancer types. If a drug pair was reported to have any of these DDI events, it was classified as ADR positive or else unknown (inferring not adverse). At this stage, the synergy level and the ADR status were to categorize the drug combinations as effective or adverse. The drug pairs with synergy levels greater than or equal to +3 and no reported adverse DDI were classified as effective while those with synergy levels less than or equal to -2 and with reported adverse DDI were classified as adverse. Table 1 reports the number of effective and adverse drug combinations selected for each cancer type. These selected drug combinations were used to assess the goodness of the network distance-based classification systems for prioritising drug combinations.

**Table 1:**
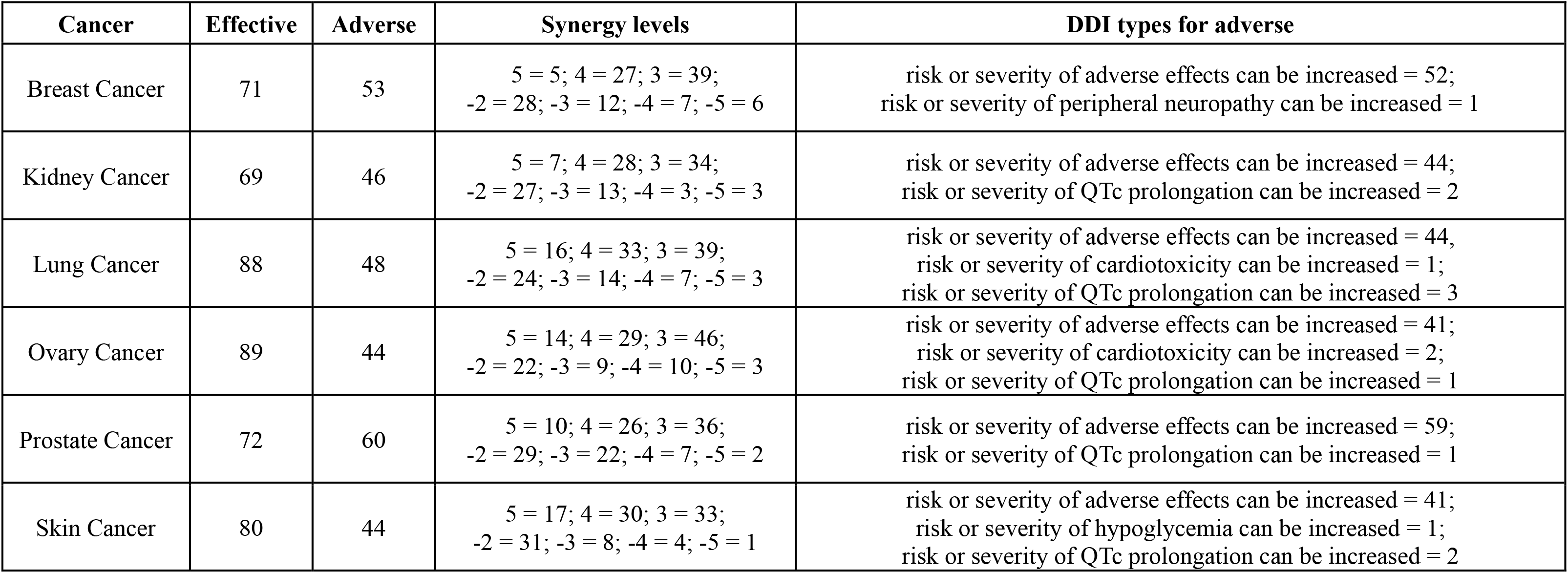
Number of drug combinations in the training data. The table also provides the distribution of the drug combination counts across different synergy levels which formed the basis for classifying the drug combinations as effective and adverse. Also listed are the different drug-drug interaction types for the adverse drug combinations.

### 2.3. Retrieval of drug targets

The targets of the drugs involved in forming the combinations were retrieved from the DrugBank database (version 5.1.10) (Wishart et al., 2018). DrugBank reports both intended and unintended targets of the drugs. More specifically, DrugBank identifies a molecule, including both proteins and nucleic acids from humans or other organisms, as a target if it is involved in the transport, delivery or activation of the drug (Wishart et al., 2008). The drug targets are primarily identified by a literature search in PubMed (Wishart et al., 2006). They are additionally confirmed based on information from other databases like FDA labels, RxList, PharmGKB and Therapeutic Target Database (TTD) which in turn reports drug-targets associations based on manufacturer-submitted information, clinical reports, pharmacogenomics etc. In our study, only the human protein targets were retained for analysis.

### 2.4. Preparation of protein-protein interaction network

A PPI network from the STRING (version 12.0) database (Szklarczyk et al., 2021) was used to study the relationship between drug targets, disease genes and safety-related genes. In STRING, each PPI is annotated with seven ‘scores’, each determined based on different types of information from three different contexts – genomic context, functional genomic experiments or direct lab assays and prior knowledge. In our study, all possible interactions between human proteins in the database were retrieved and filtered to retain those with a combined score greater than 400 (equivalent to 0.4 in scaled combined score).

### 2.5. Gene sets for measuring proximity between drug targets and disease or safety-related genes

The disease-related gene sets were retrieved from different databases like DisGeNET (release May 2020) (Piñero et al., 2020), Intogen (release 31-05-2023) (Martínez-Jiménez et al., 2020), Open Targets (v23.09) (Ochoa et al., 2021) and PharmGKB (Whirl-Carrillo et al., 2021). In addition, the GEO signatures from the Enrichr (Kuleshov et al., 2016) database and the top 50 genes prioritized by TheTA (Failli et al., 2020), a tool to identify novel disease-associated genes, for cancer were also compiled. Only the curated gene-disease associations from DisGeNET and direct associations based on human genetics data, literature, and RNA expression profiles from Open Targets were included. Since these databases do not always include gene-disease associations for the primary cancer, the genes associated with the cancer sub-types were included. The specific gene sets were selected from the databases such as to include the tissue name e.g. “breast” and terms “cancer” or “carcinoma” or “sarcoma” in their names while excluding gene sets containing the terms “hereditary” or “familial” or “susceptibility” or “predisposition”. Notably, multiple different sources were included to avoid any specific biases due to the gene-disease association curation methods. Moreover, some of these data sources focus on specific molecular information: for example, DisGeNET focuses on genetic information, while GEO signatures are based on transcriptomic data. This inclusion thus enables the identification of key molecular aspects relevant to distinguishing effective from adverse drug combinations. Similarly, the adverse-effect-related gene sets were retrieved from databases like ADReCS-Target (Huang et al., 2018), Comparative Toxicogenomics Database (CTD) (release November 2023) (Davis et al., 2023), and DisGeNET (release May 2020). These gene sets were identified based on six adverse reaction terms – namely hepatotoxicity, immune-related reaction, cardiotoxicity, neurotoxicity, haematological toxicity, and carcinogenicity – reported in the literature to be frequently associated with post-marketing drug withdrawal (Aronson, 2017). Additional gene sets containing terms like “toxicity” and/or “drug” in their names were also extracted and compiled as safety-related gene sets. Supplementary Figure 1 shows the size of each gene set, highlighting that some may have many associated genes, while others are more specific. This helps assess if gene set size impacts network-based estimation of efficacy or safety.

### 2.6. Estimation of efficacy and safety of drug combinations

The framework uses a combination of RWR (Tong et al., 2006) and FGSEA (Korotkevich et al., 2016), an approach we previously used for addressing different computational challenges (Rintala et al., 2021; Sakhteman et al., 2021), to quantify efficacy and safety. In theory, RWR explores the global network structure and calculates the proximity of a set of given ‘seed’ nodes to all nodes in the network. These proximities, also referred to as visiting probabilities (higher is closer), can be used to rank the nodes for subsequent use as input for FGSEA to evaluate their significance in the context of predefined gene sets. In our study, drug targets of each drug combination were used as seed nodes to run the RWR algorithm as implemented in the R package dNet (version 1.1.7) with a restart probability of 0.5. A threshold was then selected using the elbow method from the resultant probabilities and genes/proteins exceeding the threshold used to perform FGSEA using the fgsea (version 1.24.0) R package. To explain in more detail, the non-seed node probabilities were plotted in descending order, followed by identifying the reference line connecting the plot’s largest and smallest probability points. The perpendicular distances of all the probability points to the identified reference line were evaluated and the probability corresponding to the highest perpendicular distance was considered as the threshold. The FGSEA was executed against a combined set of disease and adverse effect-related gene sets with minSize = 5, maxSize = 500 and scoreType = “pos” as the parameters. The obtained NES was considered the efficacy and safety estimate of the drug combinations.

### 2.7. Calculation of direct proximity

In our study, the estimates from the RWR-FGSEA approach were compared to a baseline shortest path-based approach to verify if the former would outperform the latter in distinguishing effective from adverse drug combinations. While different network-based proximity measures exist, the method proposed by Cheng et al., 2019 called separation distance was chosen as it has been shown to outperform other similar methods.

### 2.8. A classification system for anti-cancer drug combinations using network medicine and gene sets linked to disease physiology and adverse drug reactions

This section describes the classification system for distinguishing effective from adverse anti-cancer drug combinations. Both statistical and machine learning-based analyses are employed to explore if the efficacy and safety estimates derived from the union of known targets of two drugs are potentially good predictors of effective and adverse drug combinations.

*Assessing the difference between efficacy and safety estimates:* The Wilcoxon test was used to assess if estimates (or NES) from individual gene sets are sufficient to classify the drug combinations. Through a one-sided test, we specifically checked if the NES for effective drug combinations is greater in disease-related features and NES for adverse drug combinations is greater in adverse-effect features. This is based on our assumption that effective drug combinations should have higher activation of disease-related gene sets as compared to adverse-effect-related gene sets and vice versa. The gene sets showing statistical significance (at p <= 0.001) were selected for the subsequent machine learning analysis. The Wilcoxon test was also used to check the informativeness of the separation distances for distinguishing the drug combinations. However, in this case, a two-sided test was used as ideally the targets of effective drug combinations should have a lower separation distance to the gene sets irrespective of the feature type.

*Training a robust and highly interpretable classification system for distinguishing effective from adverse drug pairs:* A hierarchical classification system was implemented for distinguishing effective from adverse drug combinations. In the first stage, for each cancer type, a logistic regression classifier is trained using the estimates of each selected feature assigning 1 to the positive class i.e., effective drug combinations and 0 to the other. This training data is then used to generate predictions using the resultant logistic regression model. The sensitivities and specificities for the complete probability range generated from the model are calculated and the one corresponding to the highest balanced accuracy is identified. The inverse logistic regression function is ultimately used to solve for the NES value that would serve as the threshold for distinguishing the drug combinations. The identified thresholds are used to assign an initial classification of the drug combinations within each feature into effective and adverse represented by the votes +1 and −1 respectively. However, a slightly different voting scheme is used for the different feature types. Drug combinations receive a +1 vote for disease-related features if their NES meets or exceeds the threshold, otherwise −1. For adverse-effect-related features, the voting is reversed. In the second stage, efficacy and safety scores are calculated as the average of votes within the disease-related and adverse-effect-related features respectively. A final score is then calculated by averaging the efficacy and safety scores. The final score determines the final classification of the drug combinations. A drug combination is labelled effective if the score is positive else it is labelled adverse. The proposed hierarchical classification system offers several advantages. The clear cut-off based on the gene set enrichment makes the system highly interpretable. Averaging predictions from multiple classifications, enhances robustness, reducing the impact of individual errors for more reliable results. Its balanced two-step integration of efficacy and safety assessments ensures comprehensive decision-making while also ensuring the influence of each feature group is normalised to mitigate biases due to an unbalanced number of features (e.g., more disease-related gene sets), leading to fairer and more accurate predictions. The final predictive systems developed in our study utilize the NESs from the complete set of effective and adverse drug combinations within each cancer type. However, we implement ten iterations of a repeated three-fold cross-validation framework to assess system variability. At each iteration, the dataset was split into three folds containing equal proportions of effective and adverse drug combinations. Two of these folds were used to train the predictive system involving the aforementioned steps of fitting a logistic regression model and identifying threshold NES for each feature. The third left-out fold was used as a test set to measure the accuracy of the predictive system. The thresholds identified from the training fold were used as the basis for initial feature-wise voting on the test set data followed by the calculation of the efficacy, safety and final score. The predictive models were evaluated using three different accuracy metrics, namely balanced accuracy, sensitivity and specificity.

### 2.9. Validation against sets of drug pairs not used for the training datasets

For prediction on novel datasets, the system directly uses the identified efficacy and safety estimate thresholds to execute the voting scheme on each feature followed by calculating the efficacy, safety and final scores to assign classes to the drug combinations. Three datasets were curated to assess the generalisability of the developed predictive systems. The first dataset (Validation Data 1) is a subset of the FIMM DrugComb portal drug combination data that did not qualify as the training set. The drug pairs with synergy level +2 and no reported adverse DDI were labelled effective while those with synergy level +2 and reported adverse DDI were labelled adverse for this dataset. Although less potent, these drug combinations may still be clinically significant. The second dataset (Validation Data 2) consisted of only adverse drug combinations to verify the model’s ability to correctly identify negative class. For this dataset, the drug pairs involving at least one known licensed anti-cancer drug and having reports of an increased risk of toxicity or adverse effects were extracted from DrugBank. The list of cancer-specific licensed anti-cancer drugs was retrieved from the Cancer Drug Database (Pantziarka et al., 2021). The third validation data (Validation Data 3) was derived from clinical trial information as curated in the Continuous Drug Combination Database (CDCDB) (release 09.04.2024) (Shtar et al., 2022). In this dataset, for each cancer type, the drug combinations that had completed phase 3 or phase 4 clinical trials were labelled effective whereas the combinations whose clinical trials were terminated or withdrawn due to toxicities were labelled adverse. In all three datasets, drug combinations involving only small molecular drugs were included. Furthermore, the drug combinations were filtered to remove those that were used in training the predictive models. The number of drug combinations for each category in these datasets is summarised in Table 2.

**Table 2:** The number of drug combinations in the validation and *de novo* data sets.

| Dataset | Breast Cancer |  | Kidney Cancer |  | Lung Cancer |  | Ovary Cancer |  | Prostate Cancer |  | Skin Cancer |  |
| --- | --- | --- | --- | --- | --- | --- | --- | --- | --- | --- | --- | --- |
|  | Adv | Eff | Adv | Eff | Adv | Eff | Adv | Eff | Adv | Eff | Adv | Eff |
| Validation Data 1 | 14 | 32 | 20 | 27 | 15 | 24 | 26 | 32 | 17 | 28 | 58 | 74 |
| Validation Data 2 | 1317 | - | 396 | - | 683 | - | 934 | - | 462 | - | 591 | - |
| Validation Data 3 | - | 97 | - | 1 | 1 | 35 | - | - | 1 | 35 | 4 | 9 |
| DeNovo | 6326 |  | 6362 |  | 6312 |  | 6351 |  | 6355 |  | 6361 |  |

### 2.10. *De novo* prediction dataset

A dataset of all possible combinations of small molecular licensed anti-cancer drugs approved by the Food and Drug Administration (FDA) and European Medicines Agency (EMA) was created for screening novel drug combinations. The individual drugs used for curating this dataset were retrieved from the Cancer Drug Database. The combinations were filtered to remove those that have been used for training the predictive model or those that have already entered into clinical trials of the specific cancer type. It was also filtered to remove drug combinations that have been reported to have increased risk of toxicities and adverse effects.

## 3. RESULTS

### 3.1. Characterization of a drug combinations search space for cancer research

We utilised the drug combinations collated in the FIMM DrugComb portal as the starting point of our analysis. The drug combinations were retrieved and processed as described in the materials and methods sections to identify a robust dataset of effective and adverse drug pairs for six cancer types, including breast, kidney, lung, ovarian, prostate, and skin cancers. These six cancers were selected as they formed the bulk of data in the portal. Table 1 shows the number of drug combinations finally shortlisted for our analysis. It must be mentioned here that the relatively small number of drug combinations can be attributed to the strict filtering we applied for identifying the drug combinations. Among all the drug combinations screened, most had synergy levels between −1 and +1 implying that they may be classified as synergistic based on one synergy model but antagonistic or neutral based on another (Supplementary Figure 2). Further, only 25% of the drug combinations within each cancer type were reported to have severe adverse drug-drug interactions. Hence we selected the drug combinations that occurred at the extreme ends of the spectrums when considering the synergy level and ADR status axes. This way we prioritize drug combinations that have been experimentally shown to be effective, while also prioritizing those with known relevant adverse drug-drug interactions as adverse. Interestingly, we observed that most drug combinations in the final dataset were specific for a particular cancer type (Supplementary Figure 3). These drug pairs were mostly formed by combining kinase inhibitors with taxanes. Also, most individual drugs that participated in forming the combinations in the final dataset were distributed among both effective drug combinations and adverse drug combinations (Supplementary Figure 4). Notably, adverse drug combinations in the final dataset were mostly associated with increased risk or severity of adverse effects without the mention of the specific adverse effect that the drug combination might cause.

### 3.2. Estimating the efficacy and safety of anti-cancer drug combinations using RWR-FGSEA as a network proximity measure

The RWR-FGSEA pipeline was executed on the effective and adverse drug combinations shortlisted in the previous section. Based on initial analysis, we observed that the mean NES for effective drug combinations was higher for the efficacy-related gene sets while the mean NES for adverse drug combinations was higher for the safety-related gene sets (Supplementary Figure 5). This was also statistically confirmed by Wilcoxon’s test. In line with our expectations, we observed that there was at least one efficacy-related gene set and one safety-related gene set in each cancer type that showed statistically significant differences (at p <= 0.001) in the distribution of the NES (Figure 3). However, there were a few exceptions to the pattern. Instead of the adverse drug combinations, the effective drug combinations showed higher enrichment for the cardiotoxicity and the neurotoxicity gene sets. This result might be due to their similarity to efficacy or disease-related gene sets. Notably, the effective drug combinations had no reported adverse drug-drug interactions which necessarily does not infer that there would be no adverse reactions due to the long-term usage of the drug combinations. Furthermore, we observed that the safety-related gene sets Adverse reaction (08.06.01.018) [ADReCS], and Chemical and Drug Induced Liver Injury (MESH:D056486) [CTD] were common for all the cancer types (Supplementary File 2). Apart from these, the geneset for Drug toxicity (C0013221) [DisGeNET] also showed statistically significant differences in the distribution of the estimates. The frequent occurrence of two non-specific terms is a reflection of the adverse drug combinations included in our study i.e., they were not associated with any specific adverse reactions. This highlights the importance of the inclusion of adverse drug combinations with specific toxicities for robust identification of effective and adverse drug combinations. In contrast to safety-related features, no single source was favoured for the efficacy-related gene set. However, the genesets sourced from Intogen were significant in five of the six cancers in the study while the genesets were from OpenTargets (genetic association and somatic mutation) and DisGeNet (curated) were selected as significant in four cancers each. It must also be noted that not all the sources were represented in all cancers. For example, there were no gene sets from Intogen and OpenTargets (somatic mutation) for ovary and kidney cancer respectively. Despite this, it can be observed that disease-related genesets based on genetic associations are prioritised for the separation of effective from adverse drug combinations.

**Figure 3:**
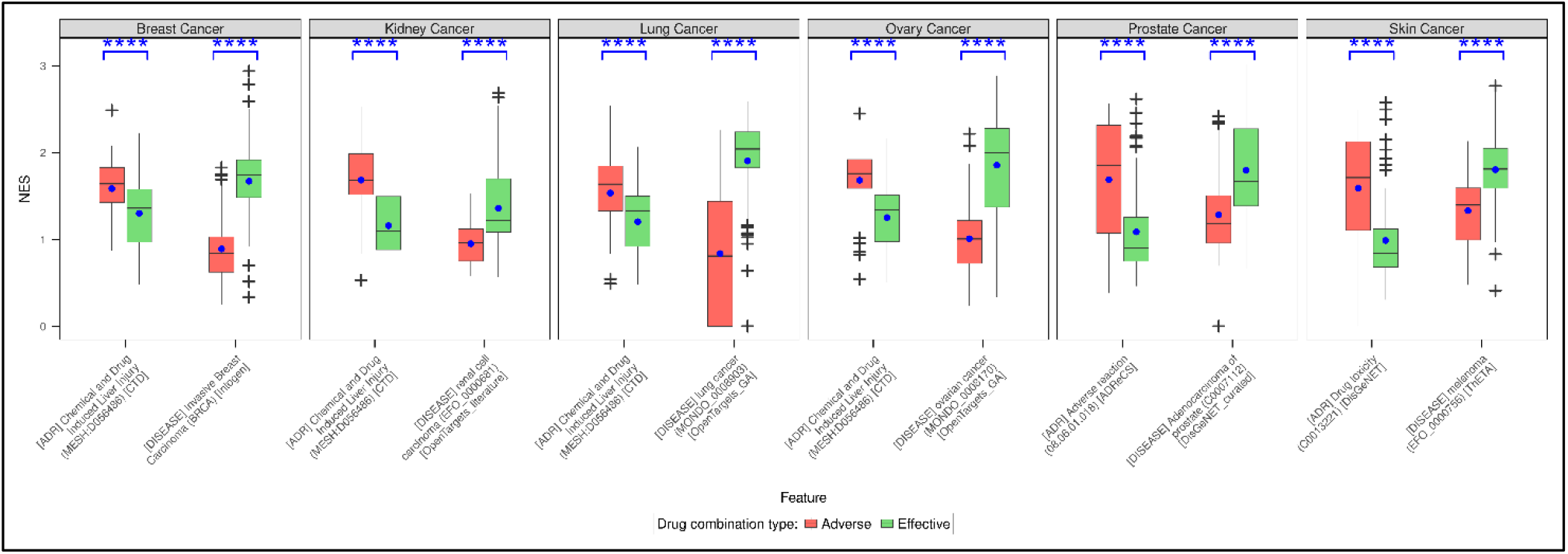
Distribution of the efficacy and safety estimates of effective and adverse drug combinations. The efficacy and safety estimates of the two groups of drug combinations obtained from the RWR-FGSEA step were compared using Wilcoxon’s test to check if they are informative in distinguishing effective from adverse drug combinations. The features that were statistically significant in the statistical test were used to develop the predictive system. Shown here are one efficacy feature and one safety feature from each cancer type that passed the statistical test and had the least p-value within each feature type. The blue dot within each box shows the mean value. The asterisks on top of each pair of boxes indicate the level of significance (***: p <= 0.001; ****: p <= 0.0001).

### 3.3. Comparison of efficacy and safety estimates with a direct network-based proximity method

We observed that the separation distances of the effective drug combination targets to either efficacy or safety-related gene sets were consistently shorter compared to the distance from adverse drug combination targets (Supplementary Figure 6). This was in line with previous findings that targets of effective drug combinations must overlap with the disease-related genes. Wilcoxon test showed that only kidney and ovary cancers had at least one efficacy and one safety feature for which the distribution of the separation distances was significantly different (at p <= 0.001) (Figure 4). In either case, the cardiotoxicity gene set from DisGeNET (C0876994) was selected among the safety features. The only other safety-related feature observed to be significant was the Chemically-Induced Liver Toxicity (C4279912) geneset from DisGeNET in kidney cancer (Supplementary File 3). In the case of lung, prostate and skin cancers, only efficacy features had significantly different separation distances. There were no features identified for breast cancer at the set threshold. Based on these observations, it can be hypothesized that the targets of both effective and adverse drug combinations are at similar proximities to safety-related gene sets and thus may not be practical enough in distinguishing the drug combinations. The proximity to efficacy-related genesets might be sufficient but it might be more appropriate to quantify how effective a drug combination would be – lower proximity inferring higher effectiveness. Unlike the direct network-based proximity method, RWR- FGSEA suggests that drug combinations must satisfy two criteria simultaneously to be classified as effective or adverse. Specifically, the effective drug combinations must have a higher efficacy estimate but a lower safety estimate, while the adverse drug combinations must have a higher safety estimate and a lower efficacy estimate. Ideally, the gap between these two estimates could dictate how effective a drug combination would be without causing any side effects.

**Figure 4:**
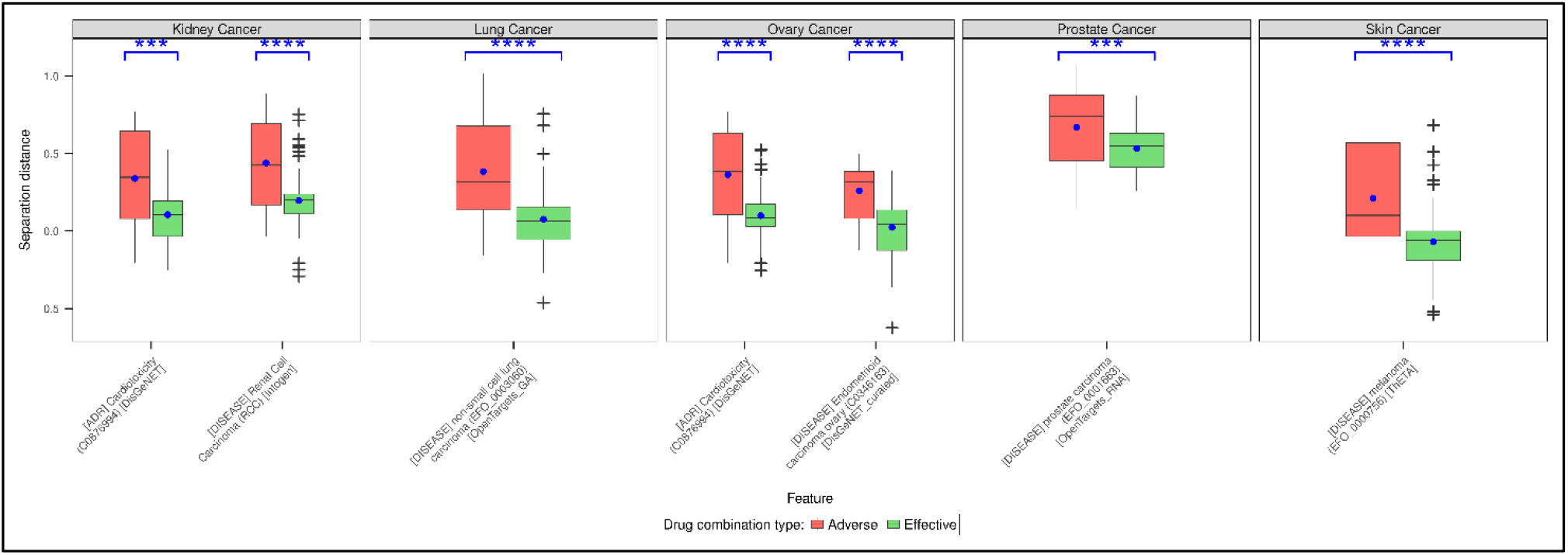
Distribution of the separation distances (proximity) of the targets of the effective and adverse drug combinations to the efficacy and safety gene sets (features). Wilcoxon’s test was also used to assess the informativeness of the separation distance of the drug combination targets to the disease-related and adverse-effect-related genes in distinguishing effective from adverse drug combinations. Shown here are one efficacy feature and one safety feature from each cancer type that passed the statistical test and had the least p-value within each feature type. If a feature of any one type is not represented, it infers that there were no significant features. The asterisks on top of each pair of boxes indicate the level of significance (***: p <= 0.001; ****: p <= 0.0001).

### 3.4. A predictive system for distinguishing effective from adverse cancer drug combinations

We leveraged the observation about differentially enriched efficacy and safety gene sets between the effective and adverse drug combinations to develop a threshold-based system to predict labels for novel cancer drug combinations. Effectively, these thresholds (Supplementary Figure 7) provide a baseline to understand how high the NES of a drug pair against a particular gene set should be for it to be classified as effective or adverse. We observed an average threshold NES of 1.262 across all cancer and feature types. Practically, for a novel drug combination to be effective, its NES against efficacy gene sets should be higher than 1.262 while its NES against safety gene sets should be lower than 1.262. If the NES exceeds the threshold values against both efficacy and safety gene sets, it would inform that the drug combination although highly effective also has a high chance of causing adverse reactions. Ideally, it would be desirable to identify drug pairs that have higher NES against efficacy gene sets and lower NES against safety gene sets.

From the variability assessment of the predictive systems using a cross-validation framework, we observed that the threshold selected based on the complete data was either equal to or close to the median threshold identified over the performed ten iterations (Supplementary Figure 8). It must be mentioned here that predictive systems presented in the study use thresholds based on the complete dataset (Supplementary Figure 7). Further, we observed that the median balanced accuracy for both the training and test data was above 0.75 for most cancer types (Figure 5). A deviation was observed in the case of balanced accuracy on the test data for breast cancer where it remained below the 0.75 mark. Additionally, the predictive systems for breast cancer and kidney cancer had a median specificity (true negative rate) below 0.75 but were compensated by the higher sensitivity (true positive rate) during cross-validation. For prostate cancer, which had the least effective-to-adverse drug combination ratio, the situation was inverse i.e., the predictive system had a lower sensitivity and was compensated by higher specificity. On the final predictive system based on the complete dataset though, we observed that the predictive systems for all cancer types achieved a minimum balanced accuracy of 0.75. Consistent with the variability check using the cross-validation framework, we observed that even in the final predictive system, the breast cancer system had a lower specificity but was compensated by higher sensitivity while the prostate cancer system had low sensitivity and was compensated by higher specificity.

**Figure 5:**
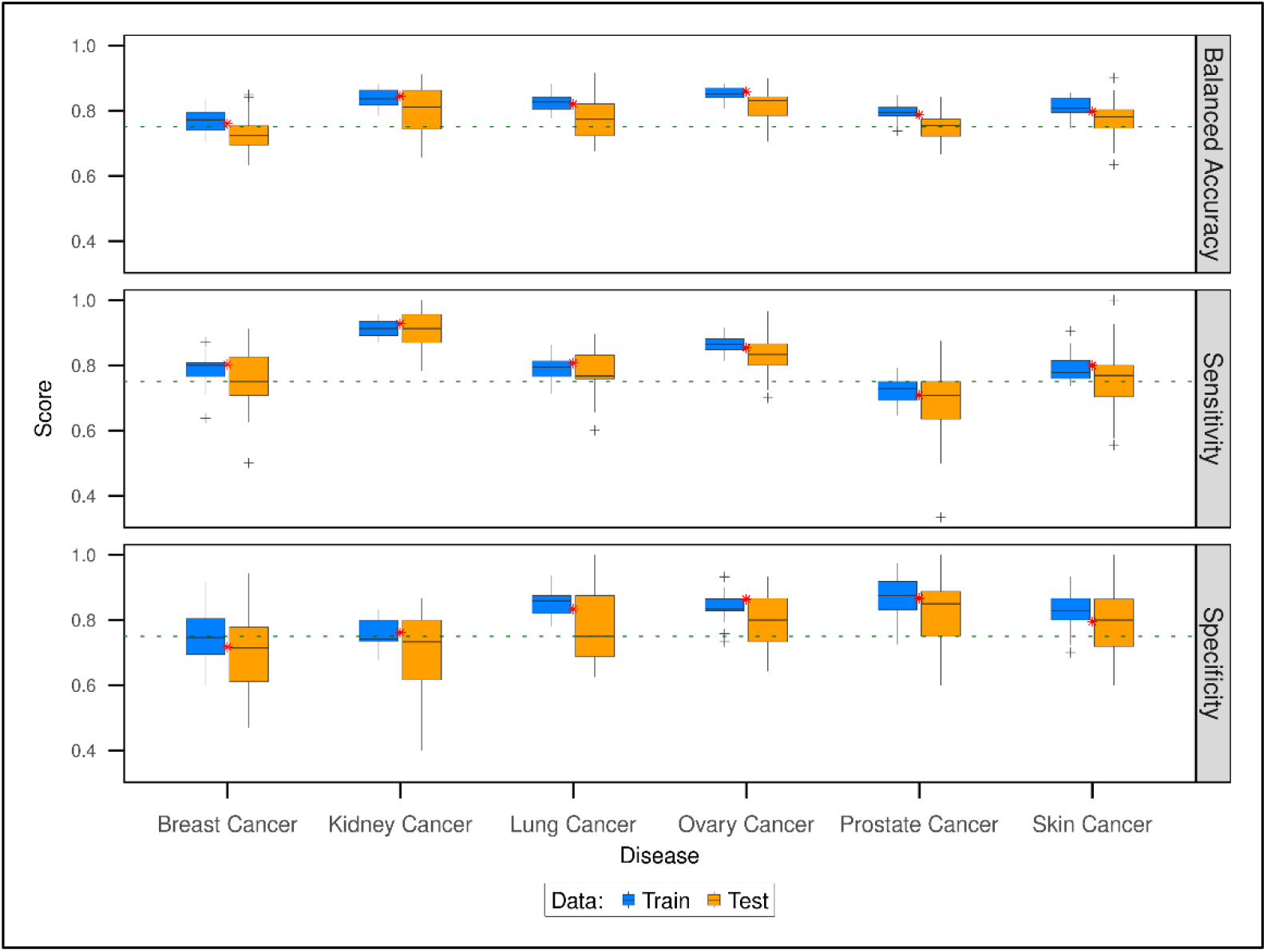
Accuracy of the predictive system. The efficacy and safety estimates obtained from the RWR-FGSEA approach were used to develop predictive systems to distinguish effective from adverse drug combinations in each cancer type. The box plots here show the accuracy scores of the systems on training and test partitions of the data during the ten iterations of three-fold cross-validation. The red asterisk shows the final accuracy score of the predictive system trained on the complete dataset.

### 3.5. Assessment of the predictive system on external datasets

The proposed predictive system for identifying and ranking cancer drug combinations was assessed against sets of untested drug combinations, labelled either effective or adverse. For the validation data 1, our predictive system showed a good positive prediction rate (mean sensitivity = 0.78) in all cancers except prostate cancer (Figure 6a). However, the negative prediction rate was low, with the kidney cancer system achieving the highest specificity of 0.55. Arguably, this might be attributed to the comparatively lower number of adverse drug combinations in the dataset. The mean balanced accuracy, a better metric for assessing imbalanced situations, was observed to be 0.61, indicating 61% accuracy in distinguishing effective from adverse synergistic drug combinations. To mitigate the impact of the small number of adverse drug combinations, we assessed the system using only negative labels i.e., only adverse drug combinations (Validation Data 2). The aim was to check the false positive rate as any drug combination falsely classified as effective and safe might lead to detrimental consequences. Here, the lung and the kidney cancer systems performed the best (specificity = 0.85) and the worst (specificity = 0.09) respectively (Figure 6b). The remaining four cancers achieved an average specificity of 0.62. A deeper inspection revealed that the overall specificity was driven by how the system predicts the dominating sub-class of drug combinations (i.e., labelled for an increase in risk or severity of adverse effects) within each dataset. Interestingly, the predictive system for breast cancer was able to predict the drug combinations leading to neurotoxicity and nephrotoxicity with high accuracy (specificities 0.96 and 0.85 respectively) even though the overall specificity was a mere 0.53. The predictive systems showed variable performance when assessed on validation data 3 (Figure 6c). The kidney and the prostate cancer systems acquired perfect sensitivity and specificity respectively but they were represented by only one drug combination each.

**Figure 6:**
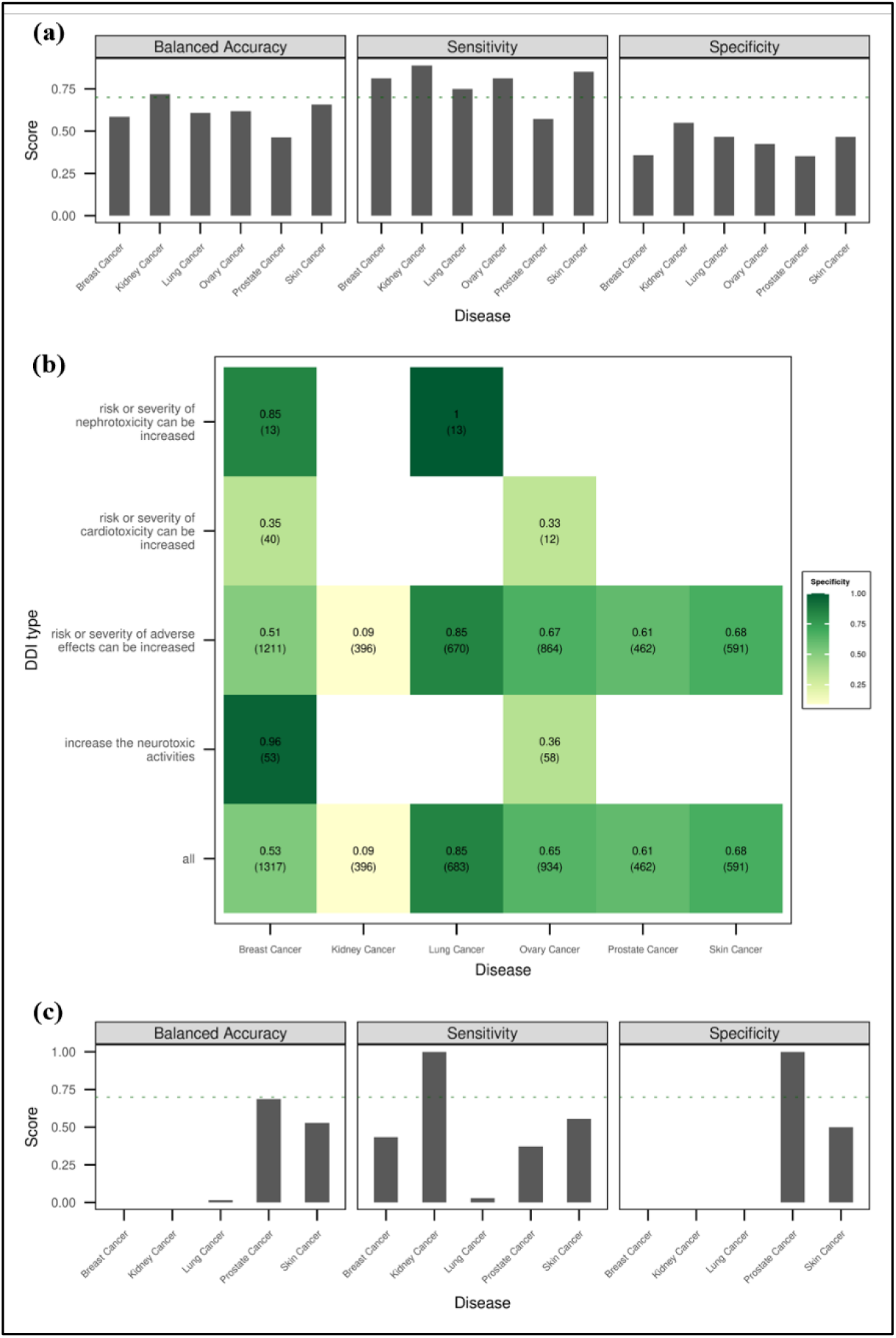
Accuracy of the predictive systems on the external validation datasets. For the validation datasets 1 and 3 (fig. a and c), sensitivity, specificity and balanced accuracy were measured. For the validation dataset 2 (fig. b), only specificity was measured as it included only negative samples.

Cumulatively, the lower accuracies of the predictive system on the external datasets might be due to the inaccuracies in the assignment of true labels for the drug combinations. For example, if a drug combination was synergistic based on only two of the five synergism scores it was labelled as effective in validation data 1. Similarly, the combinations that had completed phase 3 or 4 clinical trials were labelled effective in validation data 3. In either case, the current absence of adverse DDI reports does not completely negate the occurrence of adverse effects in the long term. For the adverse drug combinations, we included the drug combinations whose trials were terminated or withdrawn due to toxicities but the exact toxicities were unclear in several instances. Ideally, the criteria for assigning true labels in the validation data should match the training data. However, the limited data restricted us. Yet, exploring these varied assessment datasets highlighted where our method excelled and where improvements are required.

### 3.6. Identification of *de novo* drug combinations

We used our predictive system to identify novel effective and safe drug combinations that could be considered for further experimental validation. The top three selected drug combinations for each cancer type along with their mechanisms of action (MoA) (Corsello et al., 2017) are listed in Table 3. Each of these selected drug combinations received a perfect score of +1 implying that they were predicted as effective based on all the features included in the predictive system. However, as discussed in the earlier sections, our predictive systems currently do not utilise certain toxicity terms like cardiotoxicity and neurotoxicity for prediction. This can be circumnavigated by directly exploring the efficacy and safety estimates for the drug combinations. For instance, we can prioritise the drug combinations that received a perfect score of +1 based on the difference in the maximum score of efficacy gene sets and the maximum score of safety gene sets. This way we ensure that the drug combination has a higher efficacy when compared to the chances of such toxicities. The detailed scores of the top ten drug combinations predicted by our systems are given in Supplementary File 4. Among the best three-drug combinations selected to be prioritised by our system, the combination involving Panobinostat (DB06603) and Sotorasib (DB15569) was selected in three cancer types – breast cancer, kidney cancer, and skin cancer. Panobinostat is a non-specific histone deacetylase inhibitor that is used for the treatment of multiple myeloma in combination with other anti-neoplastic drugs. While Sotorasib is a KRAS inhibitor used for the treatment of non-small cell lung cancer. This combination might be of particular interest for further experimental validation as both these drugs were approved for use only in the last ten years. Particularly for Sotorasib, it was the first approved inhibitor for KRAS-mutant proteins that have been long held as one of the root causes of cancer (Blair, 2021). Another interesting drug that occurred in the priority combinations is Elacestrant (DB06374). Elacestrant is an estrogen receptor antagonist approved by the FDA in 2023. It was prioritised to be combined with Selinexor (DB11942), Ivosidenib (DB14568) and Panobinostat (DB06603) in breast, lung and skin cancers respectively.

**Table 3:** The list of selected (top three) novel drug combinations to prioritise in the six cancer types. The MoA of the drugs have been retrieved from the Drug Repurposing Hub. Additional details about these drug combinations can be found in Supplementary File 4.

| Drug1 DrugBank ID | Drug2 DrugBank ID | Drug1 name | Drug2 name | Drug1 MoA | Drug2 MoA |
| --- | --- | --- | --- | --- | --- |
| **Breast Cancer** |  |  |  |  |  |
| DB06374 | DB11942 | Elacestrant | Selinexor | Estrogen receptor degrader | Exportin antagonist |
| DB00947 | DB11942 | Fulvestrant | Selinexor | Estrogen receptor antagonist | Exportin antagonist |
| DB06603 | DB15569 | Panobinostat | Sotorasib | HDAC inhibitor | KRAS inhibitor |
| **Kidney Cancer** |  |  |  |  |  |
| DB12015 | DB14568 | Alpelisib | Ivosidenib | PI3K inhibitor | Isocitrate dehydrogenase inhibitor |
| DB06603 | DB15569 | Panobinostat | Sotorasib | HDAC inhibitor | KRAS inhibitor |
| DB11942 | DB15569 | Selinexor | Sotorasib | Exportin antagonist | KRAS inhibitor |
| **Lung Cancer** |  |  |  |  |  |
| DB00947 | DB14568 | Fulvestrant | Ivosidenib | Estrogen receptor antagonist | Isocitrate dehydrogenase inhibitor |
| DB06374 | DB14568 | Elacestrant | Ivosidenib | Estrogen receptor degrader | Isocitrate dehydrogenase inhibitor |
| DB12015 | DB15569 | Alpelisib | Sotorasib | PI3K inhibitor | KRAS inhibitor |
| **Ovary Cancer** |  |  |  |  |  |
| DB09330 | DB06603 | Osimertinib | Panobinostat | EGFR inhibitor | HDAC inhibitor |
| DB00317 | DB06603 | Gefitinib | Panobinostat | EGFR inhibitor | HDAC inhibitor |
| DB00631 | DB01259 | Clofarabine | Lapatinib | Ribonucleotide reductase inhibitor | EGFR inhibitor |
| **Prostate Cancer** |  |  |  |  |  |
| DB08899 | DB15569 | Enzalutamide | Sotorasib | Androgen receptor antagonist | KRAS inhibitor |
| DB06772 | DB08899 | Cabazitaxel | Enzalutamide | Microtubule inhibitor | Androgen receptor antagonist |
| DB11901 | DB06772 | Apalutamide | Cabazitaxel | Androgen receptor antagonist | Microtubule inhibitor |
| **Skin Cancer** |  |  |  |  |  |
| DB06603 | DB15569 | Panobinostat | Sotorasib | HDAC inhibitor | KRAS inhibitor |
| DB06374 | DB06603 | Elacestrant | Panobinostat | Estrogen receptor degrader | HDAC inhibitor |
| DB11963 | DB06603 | Dacomitinib | Panobinostat | EGFR inhibitor | HDAC inhibitor |

Note: The MoA of Sotorasib is currently not reported in the Drug Repurposing Hub. It has been filled based on DrugBank.

## 4. CONCLUSIONS

In this work, we developed a computational tool using a network medicine framework to identify and prioritise effective and safe anti-cancer drug combinations. Our proof-of-concept evaluates the mechanistic efficacy and safety of drug pairs through a combination of RWR and FGSEA approaches. Using a case study around six cancer types, we demonstrate the applicability of our proposed system for identifying effective and safe anti-cancer drug combinations. A key advantage of our framework is the requirement of minimal information for prediction, i.e., only the drug combination targets are required to predict if it will be effective or adverse. Secondly, RWR allows the testing of combinations involving two or more drugs using the same framework. Although this work mainly focuses on two-drug combinations, the validation dataset 3 demonstrates the possibility of estimating the efficacy and safety of three-drug combinations as well. Despite these advantages, our framework is not devoid of limitations. The accuracy of true labels in the training data is critical to developing a robust predictive system. For example, in the current setup, we are penalised because effective drug combinations in the training data are labelled based on the absence of reported adverse DDI, which may not reflect their true status. Additionally, the drug combinations that qualified to be adverse were mostly associated with an increase in adverse effects without specifying the actual effect. This restricted the correct identification of drug combinations with very specific side effects. Furthermore, though we developed separate predictive systems for the different cancer types, it does not account for the differences in patient sub-groups that may lead to differences in drug combination effect. In future versions of our framework, we aim to improve particularly on these two aspects to develop a predictive system for prioritisation of cancer sub-type specific drug combinations. With these improvements incorporated, we believe our predictive systems could have even better accuracy in predicting effective and safe anti-cancer drug combinations.

## DATA AVAILABILITY

Publicly accessible data were used in the study. The R scripts for analyses of the data are available on GitHub (https://github.com/UEFBiomedicalInformaticsLab/DrugCombination_1.git).

## ABBREVIATIONS

CDCDB: Continuous Drug Combination Database
CTD: Comparative Toxicogenomics Database
DDI: Drug-Drug Interactions
EMA: European Medicines Agency
FDA: Food and Drug Administration
FGSEA: Fast Gene Set Enrichment Analysis
HSA: Highest Single Agent
NES: Normalised Enrichment Scores
PPI: Protein-protein interaction
RWR: Random Walk with Restart
ZIP: Zero Interaction Potency

## ACKNOWLEDGEMENTS

The computational analyses were performed on servers provided by UEF Bioinformatics Center, University of Eastern Finland, Finland.

## AUTHOR CONTRIBUTIONS

AG: Data curation, Formal analysis, Software, Writing – original draft

VF: Conceptualization, Funding acquisition, Supervision, Writing – review and editing

