## Supplementary Figures for "Network-based estimation of therapeutic efficacy and adverse reaction potential for prioritisation of anti-cancer drug combinations"

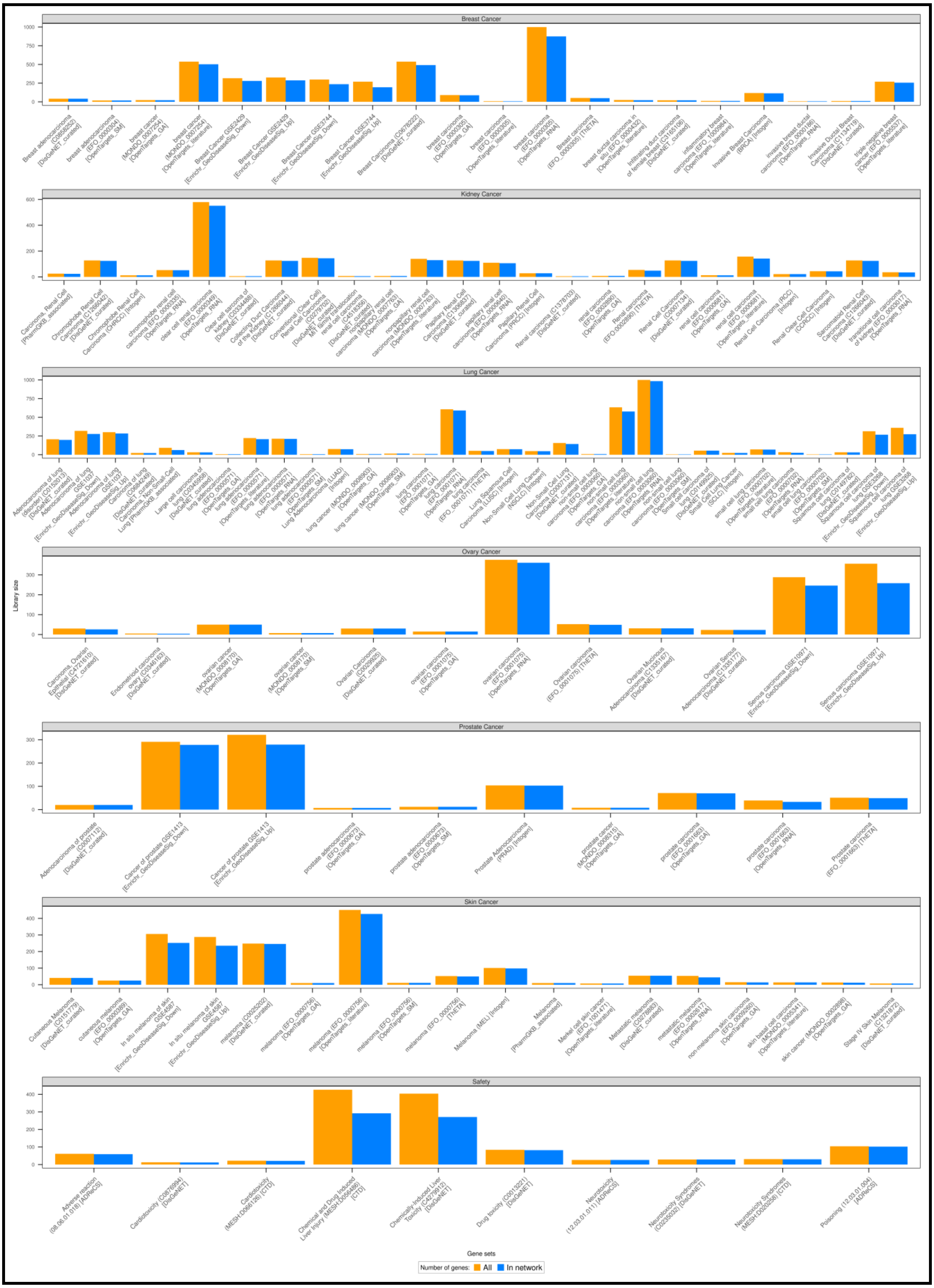


**Supplementary Figure 1:** Composition of the efficacy and safety gene set libraries. For the six cancers, we have separate gene sets for each cancer. The safety gene set is common for all cancers. The bar plot shows the number of genes in each gene set. The two bars for each gene set denote the number of genes that were compiled from the databases (orange bar) and those in the PPI network (blue bar).

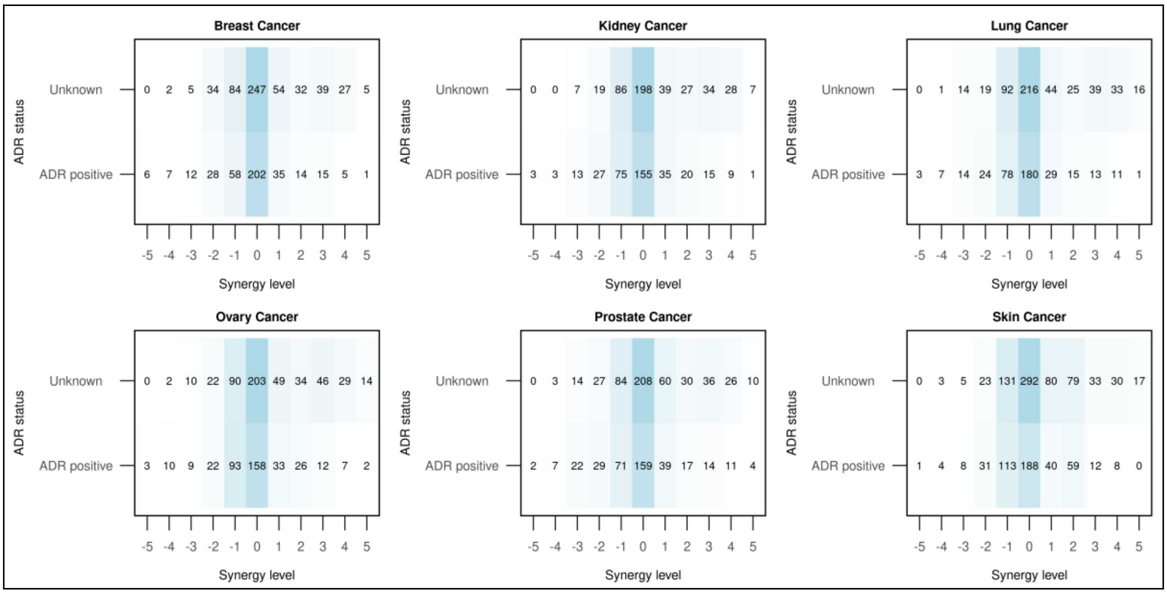


**Supplementary Figure 2:** Distribution of drug combinations across the synergy levels and ADR status. Most drug combinations had synergy levels between -1 and +1. The drug combinations with synergy level 3 or more and unknown ADR status were labelled as effective. On the other hand, the drug combinations with synergy level -2 or less and ADR positive were labelled as adverse.

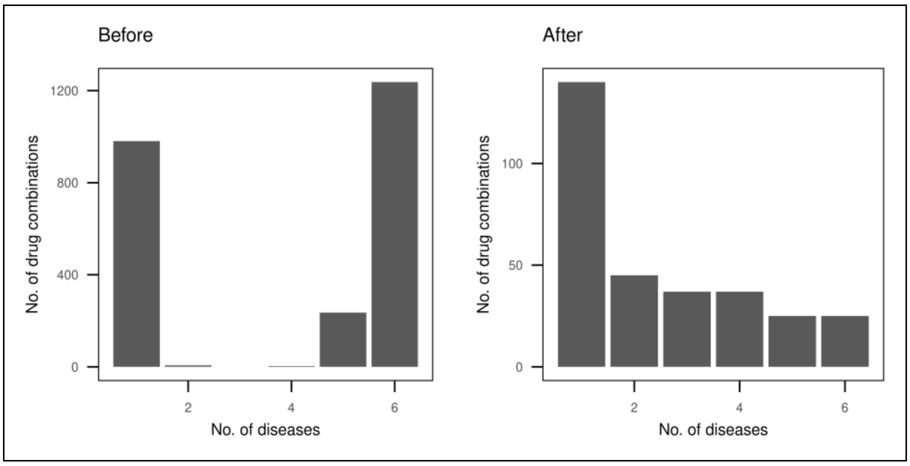


**Supplementary Figure 3:** Overview of the specificity of the drug combinations. The initial list of drug combinations identified from the DrugComb portal was mostly for use against any type of cancer. After filtering, the data set primarily included cancer-specific drug combinations.

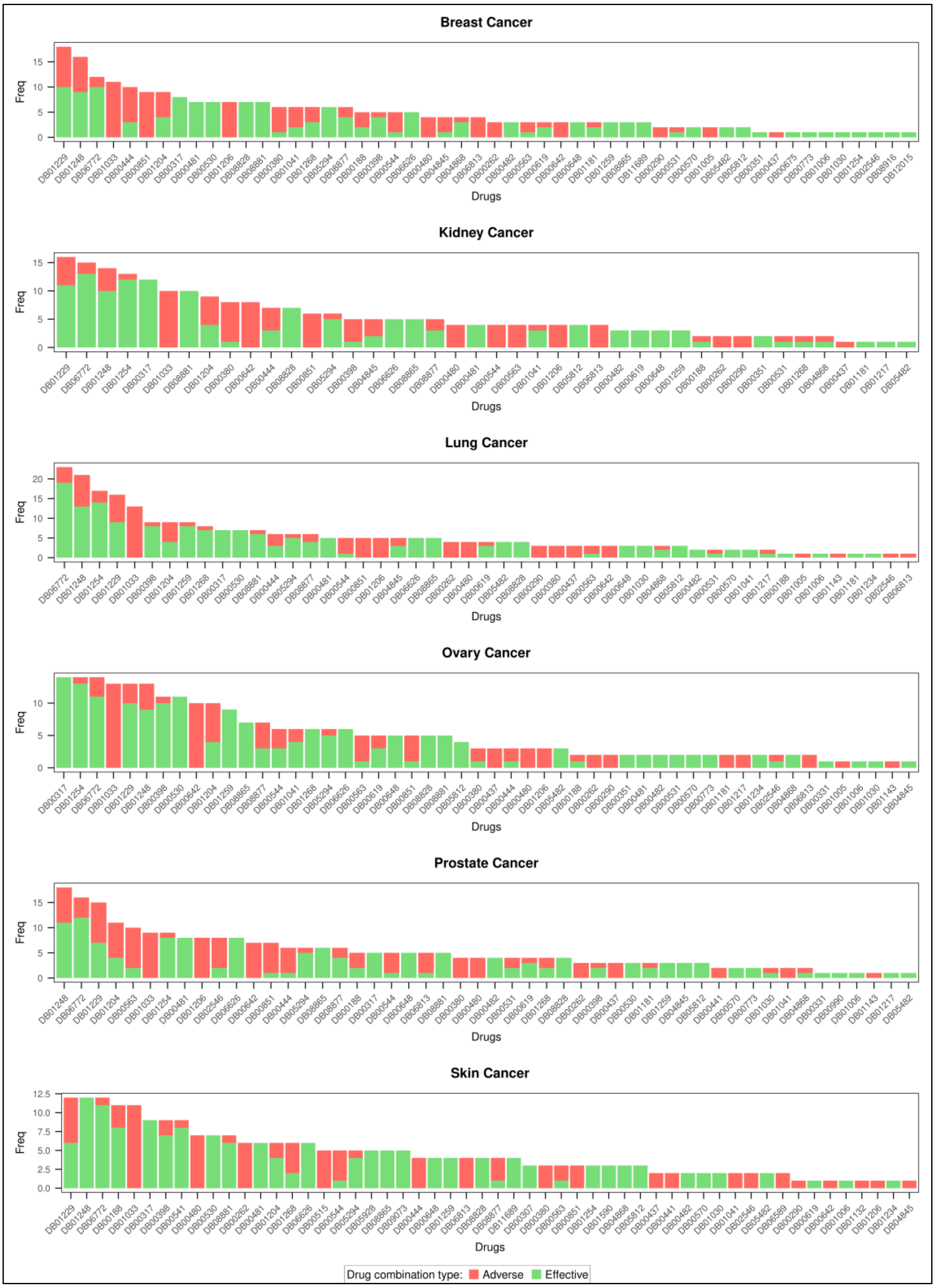


**Supplementary Figure 4:** Involvement of individual drugs in forming the drug combinations. Most individual drugs are responsible for forming both effective and adverse drug combinations. There is no absolute bias in the composition of the drug combinations i.e., one set of drugs is exclusively involved in forming the effective drug combinations while another set of drugs is involved in forming the adverse drug combinations.

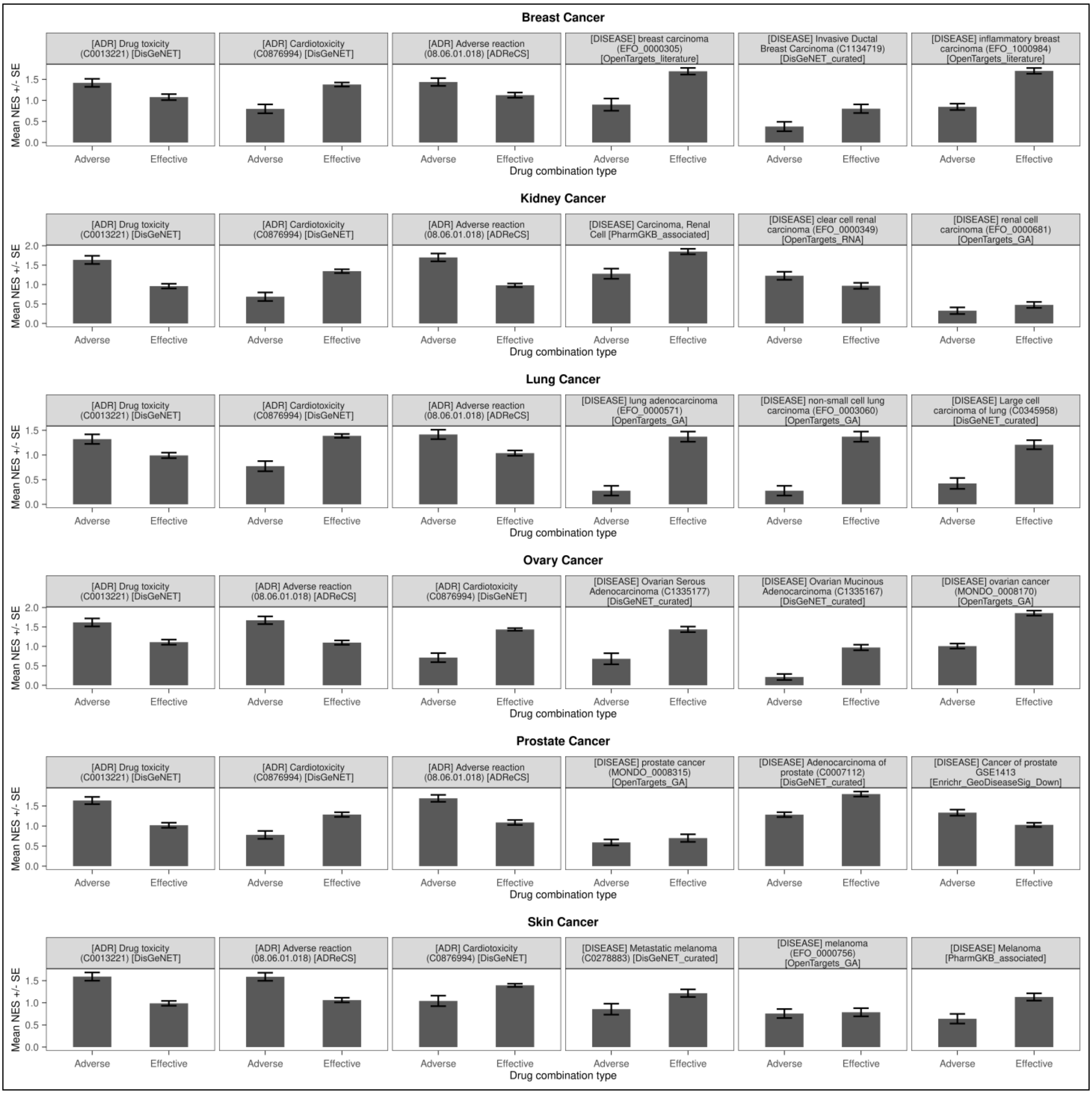


**Supplementary Figure 5:** Mean efficacy/safety estimate of the top three efficacy and safety gene sets (features) by variance in each cancer type. The safety features are tagged with '[ADR]' while the efficacy features are tagged with '[DISEASE]' at the beginning of the feature names.

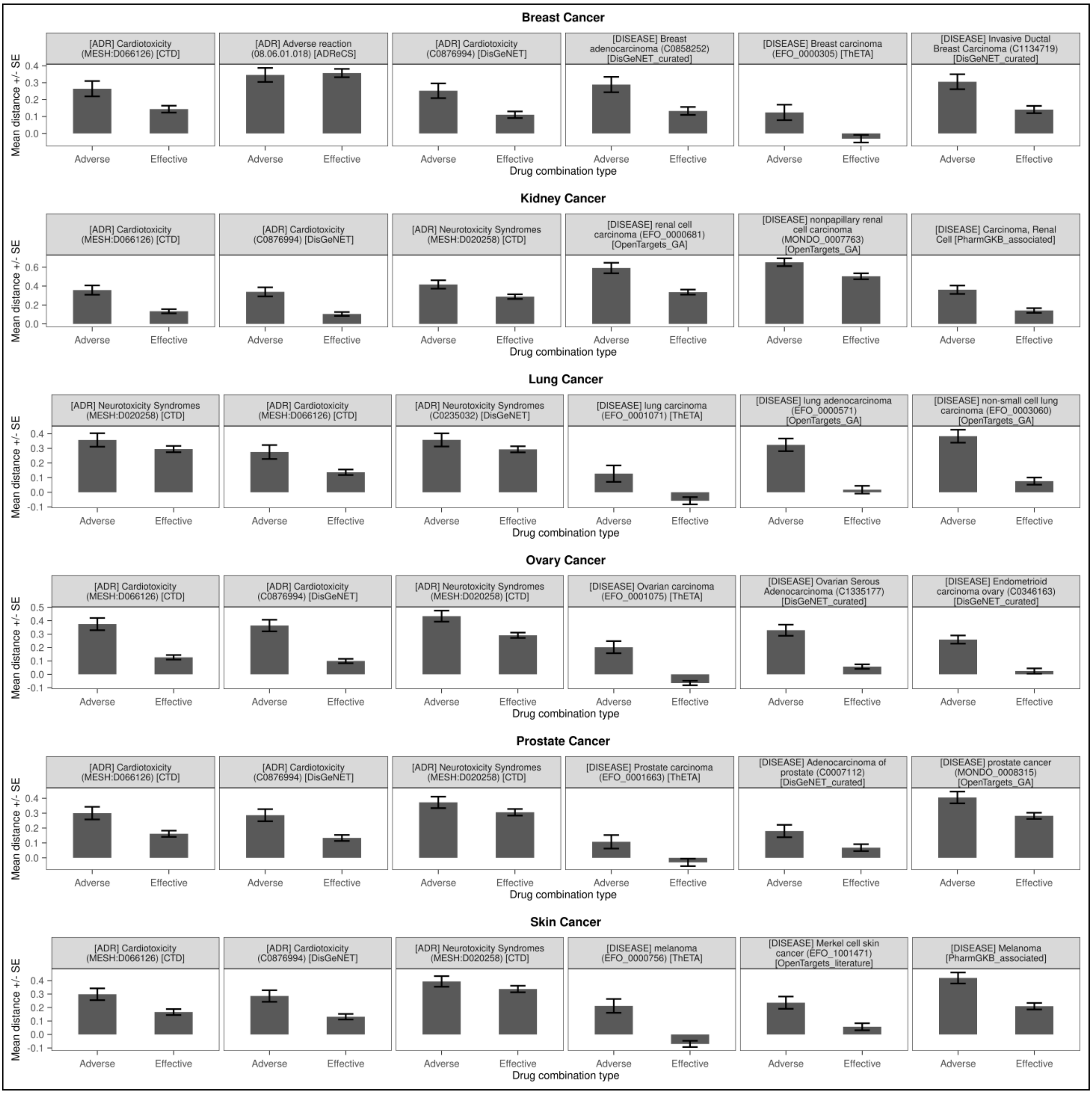


**Supplementary Figure 6:** Mean separation distance (proximity) of the top three efficacy and safety gene sets (features) by variance in each cancer type. The safety features are tagged with '[ADR]' while the efficacy features are tagged with '[DISEASE]' at the beginning of the feature names.

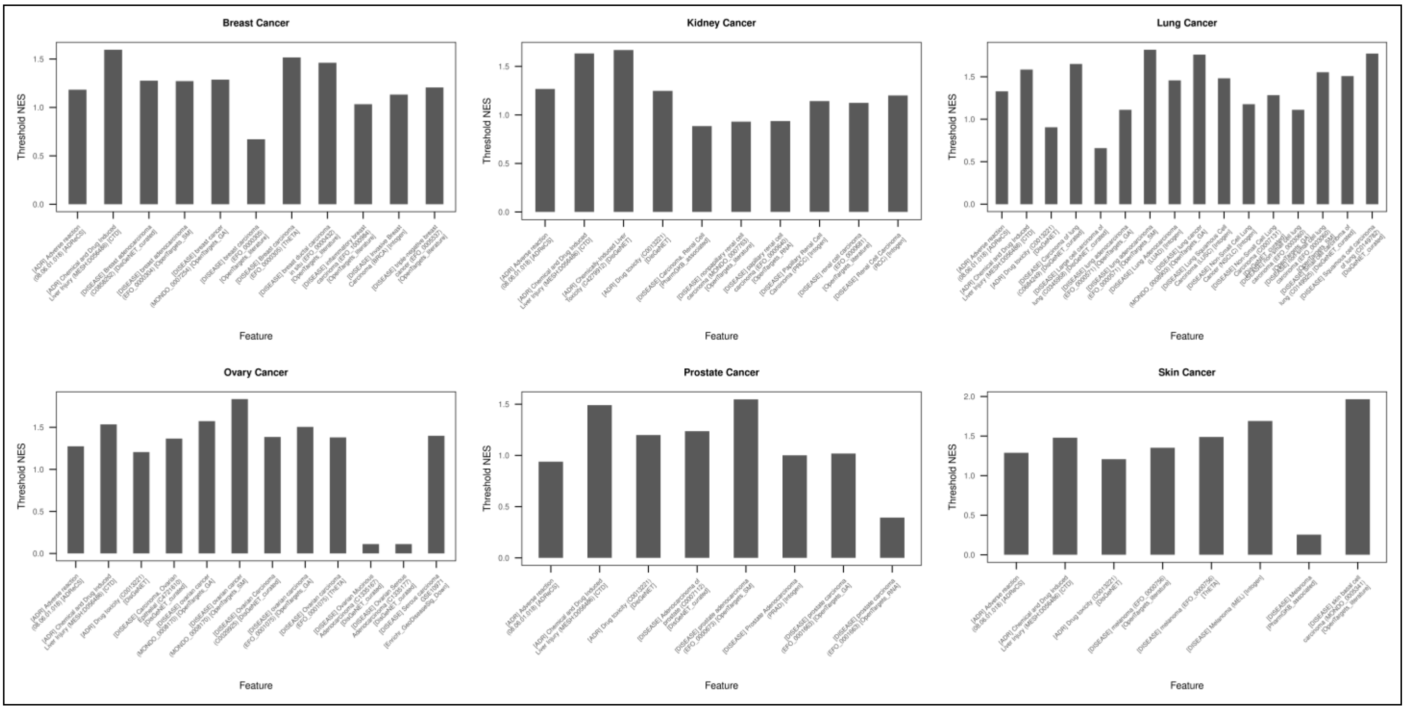


**Supplementary Figure 7:** Bar plot of the threshold estimates of the features. These thresholds form the basis of the predictive system. It defines if a drug combination will be categorised as effective or adverse based on that feature.

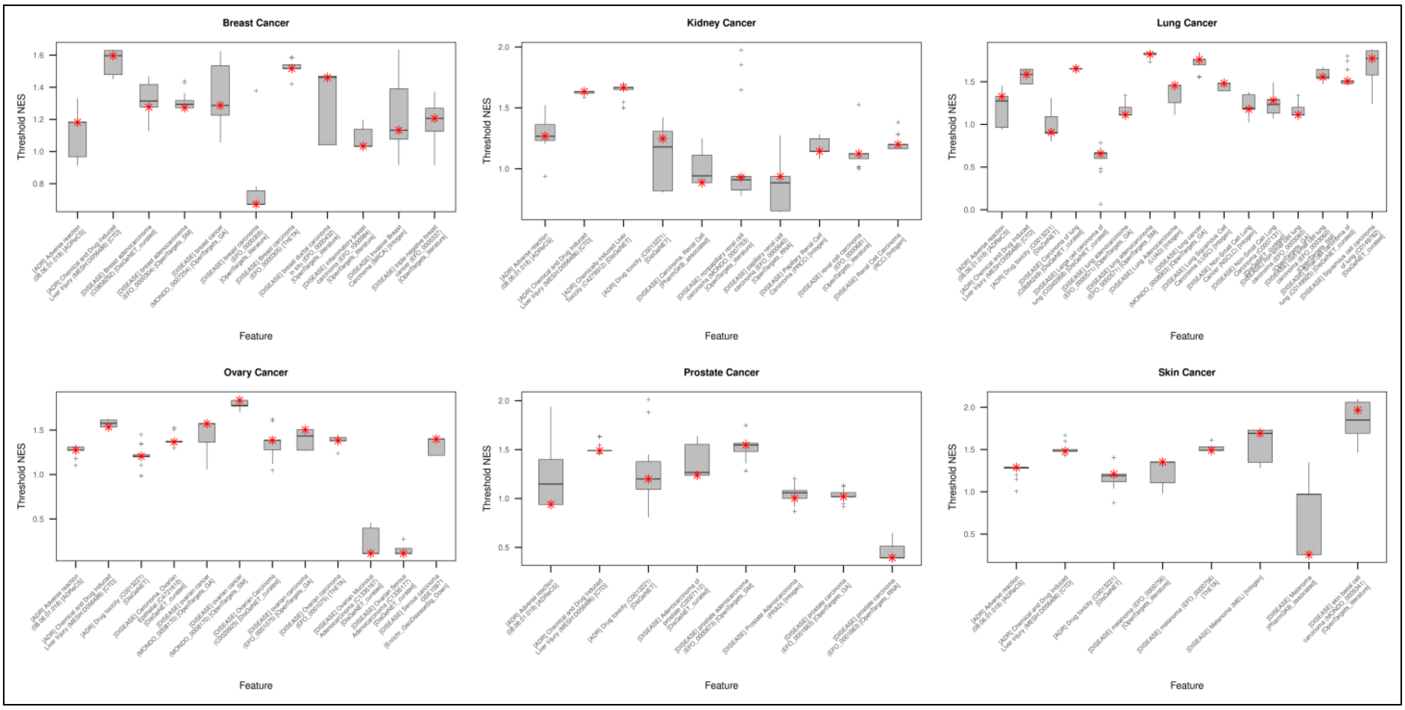


**Supplementary Figure 8:** Box plot showing the variability of the identified thresholds during the ten iterations of the three-fold cross-validation framework. The red asterisk marks the final threshold value identified based on the complete data.
